# Quantifying growth and lodging in Tef (*Eragrostis tef*) with Uncrewed Aerial Systems (UAS)

**DOI:** 10.64898/2026.01.06.697717

**Authors:** Keely E. Brown, Haley Schuhl, Dhiraj Srivastava, Getu Beyene, Mao Li, Noah Fahlgren, Katherine M. Murphy

## Abstract

Lodging is a major contributor to decreased yield in tef, a staple cereal crop in Ethiopia. Semidwarf varieties have been developed with a goal to increase yield through reduced lodging, but studying lodging susceptibility currently requires a labor-intensive, imprecise, manual scoring method. Here we present workflows for analyzing tef stand height from UAS sensors across time to both predict lodging later in the season with early height and to measure the severity of lodging after a storm event. We compare 3D point clouds generated by photogrammetry from RGB images with those generated from LiDAR to estimate height, demonstrating that they produce similar results, despite differences in cost. Stand height and lodging can both be accurately measured with low-cost UAS, reducing the need for manual measurements and increasing precision and temporal resolution in plant breeding programs.

**Significance Statement:** Extreme weather or heavy grain can cause plant stems to bend, a process called lodging. Lodging significantly reduces crop yields globally, particularly in grain crops such as tef (*Eragrostis tef*). Semidwarf crops have previously been reported to be lodging-resistant, increasing crop yields. Here, we used uncrewed aerial systems (UAS) to measure plant growth, height, and lodging in gene edited semidwarf tef lines, and compared the results to ground-truth data. Using a UAS equipped with a red-green-blue (RGB) camera or LiDAR sensor, we measured plant height and lodging, and found that early-season height measurements could predict future lodging potential. The tools used were contributed to the open-source software PlantCV-Geospatial for community use. This work contributes to a broader understanding of genetic resistance to lodging, providing valuable insights for tef crop improvement and reduces the need for labor-intensive manual measurements.

## Introduction

Tef (*Eragrostis tef* (Zucc.) Trotter) is a C4 grass generally considered tolerant to drought, flooding, and pests both in the field and during grain storage (Bekele-Alemu & Ligaba-Osena, 2023), and is a major staple cereal crop in Ethiopia (Tadele & Hibistu, 2021). In addition to favorable agronomic traits related to stress tolerance, consumer-facing traits, such as gluten-free grain with high nutritional value, are driving increasing global interest in tef (Abebe et al., 2007; Assefa et al., 2011; Gebremariam et al., 2014; Spaenij-Dekking Liesbeth et al., 2005). However, widespread adoption of tef is hindered by low yields compared to other grain crops such as maize and wheat (Mihretie et al., 2022). Lodging, competition from weeds, grain shattering, and low productivity are major limitations to tef production (Assefa et al., 2011). Like other cereals, lodging in tef is exacerbated by high wind speeds, heavy rainfall, and agronomic practices that favor top growth and often cause tef to lodge, a phenomenon in which the stem bends or snaps permanently at the base, causing the plant to fall over (Ben-Zeev et al., 2020; Berry et al., 2004; Merchuk-Ovnat et al., 2020). Lodging in tef has been reported to reduce grain yield by up to 25% and affects the quality of both grain and straw (Assefa et al., 2011; Ben-Zeev et al., 2020; Gebru et al., 2023; Zeid et al., 2012).

Semidwarf plants have been associated with lodging resistance, so plant height has been a target for traditional breeding in many crops. In wheat and rice, lodging resistant varieties were bred using spontaneous mutations, resulting in shorter, thicker stems (Dalrymple, 1985; Hedden, 2003). Maize breeding and biotechnology efforts recently developed new short-stature varieties to increase yields by reducing susceptibility to lodging (Barten et al., 2022). In cultivated barley, lodging is known to correlate with stem height and thickness (Haaning et al., 2020). Tef stand height and lodging resistance is influenced by variety, row spacing, seeding rate, and more (Blösch et al., 2020; Tasew et al., 2024; Wato, 2019). To accelerate the process of traditional breeding, Beyene et al. (2022) developed CRISPR/Cas9-based genome edited tef lines with reduced plant height and improved lodging resistance, tested in a controlled growth environment.

In order to evaluate lodging resistance and its relationship to stature for breeding pipelines, researchers must accurately measure plant height. However, the current standard methods for measuring plant height are time-consuming and manual (Sun et al., 2018; Wang et al., 2018). Additionally, to associate variation in height with lodging resistance, a lodging index must be calculated from a subjective scale and human estimations for the percentage of a plot at each lodging score (Caldicott & Nuttal, 1979). This method provides information about the overall extent of lodging across a plot, but remains subjective and time-consuming. Another challenge in studying lodging is that researchers must wait for an event like a heavy storm and return to the field to manually score soon after, which may not always be possible.

High-throughput phenotyping provides new opportunities to accurately, precisely, and efficiently quantify plant height and lodging with imaging and computational methods to measure plants and plots (Pauli et al., 2016; Shi et al., 2016; Vergara-Díaz et al., 2016; Wilke et al., 2019). Measurements made from images captured from Uncrewed Aerial Systems (UAS) and satellites are becoming increasingly popular due to their accuracy and reduced manual labor (Haghighattalab et al., 2016; Hoffmann et al., 2018; Jindo et al., 2021). In particular, drones equipped with RGB (red-green-blue), multispectral, hyperspectral, and/or LiDAR (Light Detection and Ranging) sensors have provided researchers with measurements of plant height, lodging, canopy cover, plant architecture, and more (Ayankojo et al., 2023; Barbedo, 2019; Matias et al., 2022). The reduced labor needs not only provide more reproducible measurements, but the opportunity to increase the frequency or number of measurements feasibly taken in a growing season. Thus, rather than manual lodging estimations during crop growth as well as after a weather event, lodging can be measured as the change in height before and after an event, providing a more holistic and dynamic approach to plant measurements over time.

Generating 3-Dimensional (3D) reconstructions of plants and plots has been shown to provide useful information on plant height and other measurements, but there is a trade-off between sensor cost and accuracy. LiDAR uses laser pulses to measure distances from the sensor to objects below, making it particularly useful for estimating plant height, stand height, and canopy density (Swinfield et al., 2019; ten Harkel et al., 2019). However, LiDAR sensors and the UAS required to carry their often heavy payloads are generally more expensive than RGB cameras and their associated UAS, and data analysis is often complex. RGB imaging can be used to generate 3D point clouds using structure from motion analysis methods (Leberl et al., 2010), which may have reduced accuracy compared to LiDAR (White et al., 2013). While these methods are exciting, it is critical that new software tools and methods to analyze these datatypes are accessible and reproducible so that other researchers can utilize the measurement pipelines. Lack of usable software remains a well-documented challenge in the plant phenotyping community (Lobet et al., 2013).

In this study, we compare manual measurements of plant height and lodging for a group of semidwarf tef lines and their parental control with measurements obtained using a UAS equipped with RGB and LiDAR sensors. We present reproducible, sustainably maintained analysis pipelines for estimating height using digital elevation models and the open-source software package PlantCV-Geospatial. UAS-based phenotyping generated accurate measurements of stand height and lodging from both RGB and LiDAR data. This confirmed that semidwarf tef lines were less likely to lodge during vegetative growth, in extreme weather events, or during the grain filling stage. UAS with either RGB or LiDAR sensors thus provide new opportunities for researchers and farmers to evaluate plant varieties for improved lodging resistance without subjective, manual measurements.

## Materials and Methods

### Plant growth

Tef lines were grown at the Donald Danforth Plant Science Center Field Research Site in St. Charles, Missouri, USA (38.848 N, 90.458 W) from June to September 2023. Two wild-type lines (cultivar Magna) and seven genome edited, semidwarf tef lines were planted each on a 12 m^2^ plot (4 m x 3 m) replicated three times in randomized complete block design (a total of 27 plots). Plots 101, 102, 103, 901, 902, and 903 were wild-type (Figure 1B). Plots 201, 202, and 203 were *sd-1*-1 lines, previously published (Beyene et al., 2022), and the remaining five were lines generated targeting tef orthologs of known dwarfing genes. Row spacing was 30 cm and seeds were planted in each row manually using a salt shaker to give an estimated 1-2 cm spacing between plants. The distance between plots was 2 m and between replicated blocks 3 m. Plants were irrigated with sprinkler irrigation as needed and plots were kept weed-free by manual hoeing. The field used for tef planting was preceded by soybean the previous year, and chemical fertilizers were not applied during the growing season. A severe windstorm occurred on July 29th, 2023, ∼8 weeks after planting. Wind speeds reached almost 22 mph (35.4 kph), as compared to averaging less than 4 mph (6.4 kph) the day before the storm (measured by a PheNode environmental sensor, Agrela Ecosystems, Inc.; Fig. S1).

**Figure 1.**
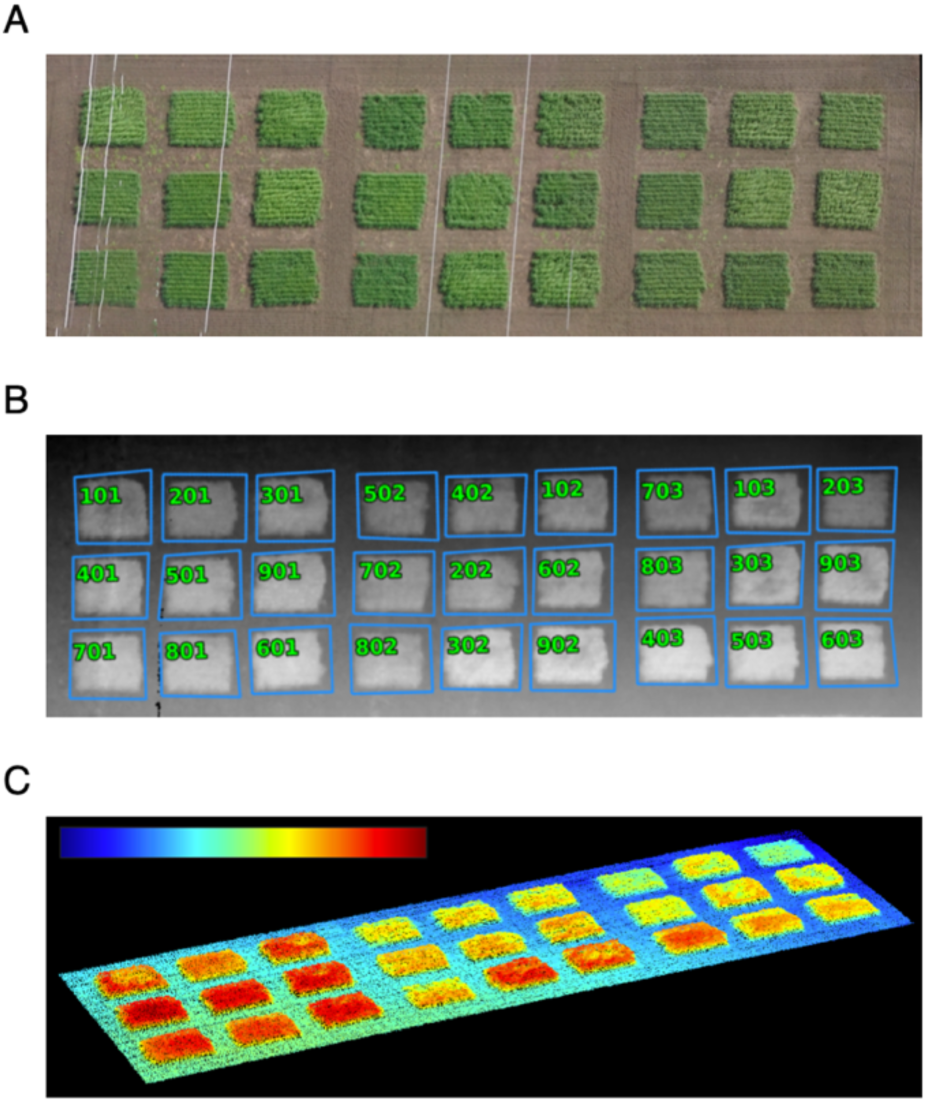
UAS orthomosaics allowed for clear plot delimitation. (A) RGB orthomosaic from July 17th, 2023. Note the powerlines (white lines) above the field. (B) DEM created from the point cloud during stitching of the orthomosaic. The powerlines were converted to no data values by PlantCV-Geospatial using a height threshold. (C) LiDAR from the same date produced point clouds for height calculation. Higher points are toward the red end of the scale.

### Manual data collection

Stand heights per plot were measured during vegetative growth at 6 weeks after planting (July 13, 2023). Stand height was measured from three different locations in each plot by measuring the height of tef plants from ground level to the top of the canopy. To do so, an A4 sized piece of paper was placed on top of standing plants in a uniform section of the canopy, and measured from the ground to the paper with a ruler. The process was repeated at three randomly selected positions in the plot, but only for standing, non-lodged plants. No plant was bent in the process, as they supported the lightweight paper. Lodging index was measured at 15 weeks after planting, which was 2 weeks after the last UAS flight, as per Caldicott and Nuttal (1979). In brief, plots were scored from 0 (no lodging) to 5 (flat plants) on a subjective severity scale, and the percent of the plot with each score was estimated to calculate a final lodging score (Caldicott & Nuttal, 1979).

### UAS data collection

UAS images were acquired at 12 timepoints throughout the growing season, roughly every 1-2 weeks from the middle of June to the end of August. Images were captured with a DJI M600 UAS mounted with RGB (Sony) or LiDAR (Phoenix) sensors, and flown at 80 m with a constant horizontal speed of 28.8 kph above the ground for all flights. Front and side imaging overlap for RGB images was 70%. Image resolution for RGB orthomosaics is reported in Table S1. For LiDAR, the pulse repetition rate was 700kHZ, the field of view was 90 degrees, with two returns per pulse and an accuracy of ∼2 cm.

### RGB image analysis

Raw images were used to construct orthomosaics using the photogrammetry software Agisoft Metashape (Agisoft LLC, St. Petersburg, Russia) through the Data to Science (D2S) platform (Jung et al., 2024; Jung, M., B. G. Hancock, Z. C. Qian, N. Zhuo, Z. Gong, J. S. Doucette, J. Jung., n.d.), which also created a digital elevation model (DEM) representing height of pixels in meters above sea level in the orthomosaic from a dense point cloud. The orthomosaic blending mode was Mosaic with “Fill Holes” enabled, the point cloud quality was set to high, the align photos accuracy was set to high, surface was DEM, and the software version of Agisoft Metashape Professional was 2.1.3 build 18946. Both orthomosaics and DEMs were georeferenced using ground control points with the QGIS Georeferencer tool (Dawson et al., 2025). Georeferencing was done to a single timepoint as a reference rather than to an absolute coordinate system. To do so, a reference orthomosaic was opened in the main map canvas, and the second orthomosaic was opened in the Georeferencer window. The August 10th flight was used as the reference orthomosaic because this was the timepoint used to create shapefile plot boundaries. Ground control point coordinates were entered by clicking on corresponding points in the two orthomosaics. Transformation type was Projective, resampling method was Nearest Neighbor, and 8 ground control points per orthomosaic were used to calculate the transformation for georeferencing. A shapefile was also made in QGIS to generate polygons around each tef plot on August 10, and used on the remaining images after georeferencing.

To measure height, georeferenced orthomosaics and DEMs were cropped to the field and opened in Python using the PlantCV-Geospatial package, which is a library of Python tools for analyzing geospatial data. PlantCV is a free, open-source image analysis package for analyzing images of plants (Gehan et al., 2017; Schuhl et al., 2025) that provides a framework for measuring and storing observations extracted per object within each image. All code associated with these analyses is available on GitHub (https://github.com/danforthcenter/teff-manuscript), as well as the PlantCV-Geospatial package (https://github.com/danforthcenter/plantcv-geospatial). As observed in the orthomosaic (Fig. 1A), tef plots were planted under power lines in the field, which could not be flown under due to UAS safety restrictions. Pixels belonging to powerlines needed to be removed to measure plot heights. During import, PlantCV-Geospatial was used with a height percentile threshold for filtering, so values above the threshold were converted to no data values. We used a threshold of 0.995 (unitless, 0-1 range), which was sufficient to remove powerlines in this field (Fig. 1B).

To obtain the stand heights of each plant pixel in the tef plots, pixel height needed to be subtracted from the elevation of the soil with respect to the mean sea level. Because the land across the field was not uniformly level, a single soil elevation could not be used for all plots. However, the soil surrounding each individual tef plot was relatively flat (Fig. 1B). Therefore, shapefiles per plot included a perimeter of soil so height was measured by the difference between plot and soil elevation (Fig. 1). For each plot, the soil elevation was estimated to be the 1st percentile in the distribution of plot height (Fig. 2), calculated using ranked pixel values from each plot’s DEM. This soil value (red) was then subtracted from the 95th percentile (blue) in the height distribution of each plot, which was estimated to represent an average canopy height (Fig. 2).

**Figure 2.**
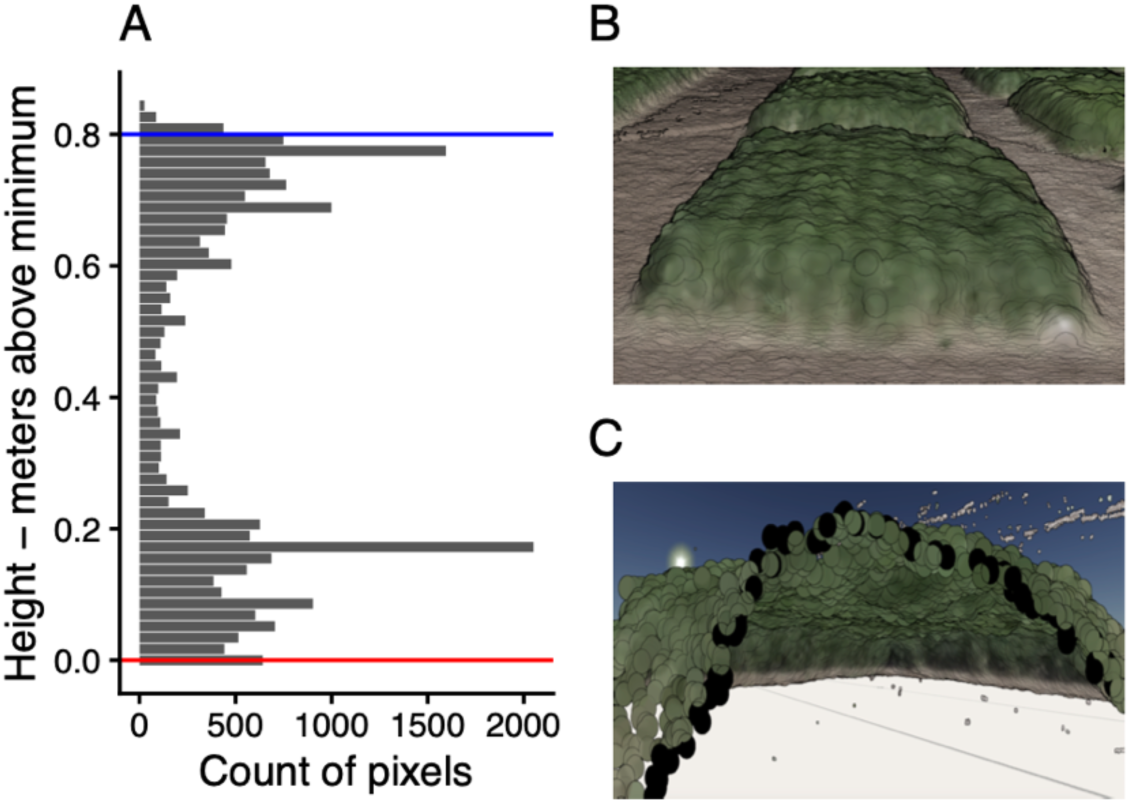
(A) Stand height was calculated as the difference between top (95%, blue line) and bottom (1%, red line) percentiles from the point cloud distribution. (B) An example of the 3D point cloud showing surrounding soil pixels. (C) A slice inside of a plot in the point cloud showing the distribution of plant heights across the canopy.

### LiDAR data analysis

The LiDAR data in LAS format was imported into MATLAB (R2024a) as a 3D point cloud. An image acquired on July 17, 2023 was selected as a reference for defining the region of interest (ROI) of the tef field. A customized algorithm was developed, allowing rotation and precise definition of the ROI boundaries by selecting the upper-left and lower-right corner points of the field. Based on prior knowledge of tef plant height ranges, a height threshold of 3 m above the ground level was applied to exclude objects and noise exceeding this height, such as power lines. Once the mask parameters were determined from the reference data, they were applied to mask all other LiDAR datasets.

The ROI was partitioned into a grid consisting of 3 rows and 9 columns (Fig. 1A). This configuration was chosen to ensure that each grid cell contained soil points. Within each grid cell, points located at least 25 cm above the minimum elevation were identified as the initial plant subset. The initial plant subsets from all grids were then combined to form the entire initial plant point cloud. A density-based noise removal method was applied to this point cloud using MATLAB’s pcdenoise() function and the refined data was segmented into clusters of interest (i.e. plots). The edge mask for each plot was generated and was subsequently applied to all ROIs in other LiDAR datasets.

We estimated the plant height using two different approaches. The first approach is similar to the RGB images, soil elevation for each plot was estimated as the 1st percentile of the plot height distribution. This soil elevation was then subtracted from the 95th percentile of the height distribution, which was used to represent the average canopy height for each plot.

The second approach is to estimate stand height by mimicking the manual measurement process, which involves placing a flat paper over the plant canopy and measuring its height above the ground. To simulate this, 10 x10 cm squares were overlaid to cover each plot, excluding the 30 cm boundary region to remove the boundary effects. Within each square, the maximum height of the plant points inside the square area were measured. The final stand height for each plot was then determined as the average height of the top 80th percentile of these maximum square heights.

### Statistics

Linear models, ANOVAs, and correlation coefficients were fit and estimated using base R (version 4.4.2) (Posit team, 2025). Linear mixed-effects models were fit using the R package nlme (Pinheiro et al., 2025) in an R environment running version 4.1.3.

## Results

### Short tef lines are resistant to lodging after a high wind event

Shorter plant varieties have consistently been shown to be resistant to plant lodging; thus, we hypothesized that semidwarf tef would both be shorter than wild-type tef grown in the field, as well as have reduced lodging. Indeed, height measured manually at a single date showed that semidwarf tef was significantly shorter than wild-type (F=12.573, p=0.00231). Semidwarf tef also had significantly lower manual lodging scores at the end of the season than wild-type (F=112.29, p=3.63e-09). Statistics are from an ANOVA including both genotype (wild-type or semidwarf) and line as factors.

### Height estimated from RGB images approximates manual measurements

We next sought to determine whether height estimated with RGB imaging would correlate to manually-collected measurements, and how additional timepoints of measurement might add temporal insights. To do so, we used the open-source software PlantCV-Geospatial to estimate height from DEMs at each date (see Methods for details). Using plant area at the 95th percentile threshold and soil area at the 1st percentile had the highest correlation to manual measurements, with a Pearson correlation coefficient (R^2^) of 0.83 (Fig. 3A), and was used as the height calculation parameters for all plots and timepoints. This supports previous research suggesting RGB imaging from UAS is an effective method for measuring stand height, as expected (Hassan et al., 2019; Matias et al., 2022).

**Figure 3.**
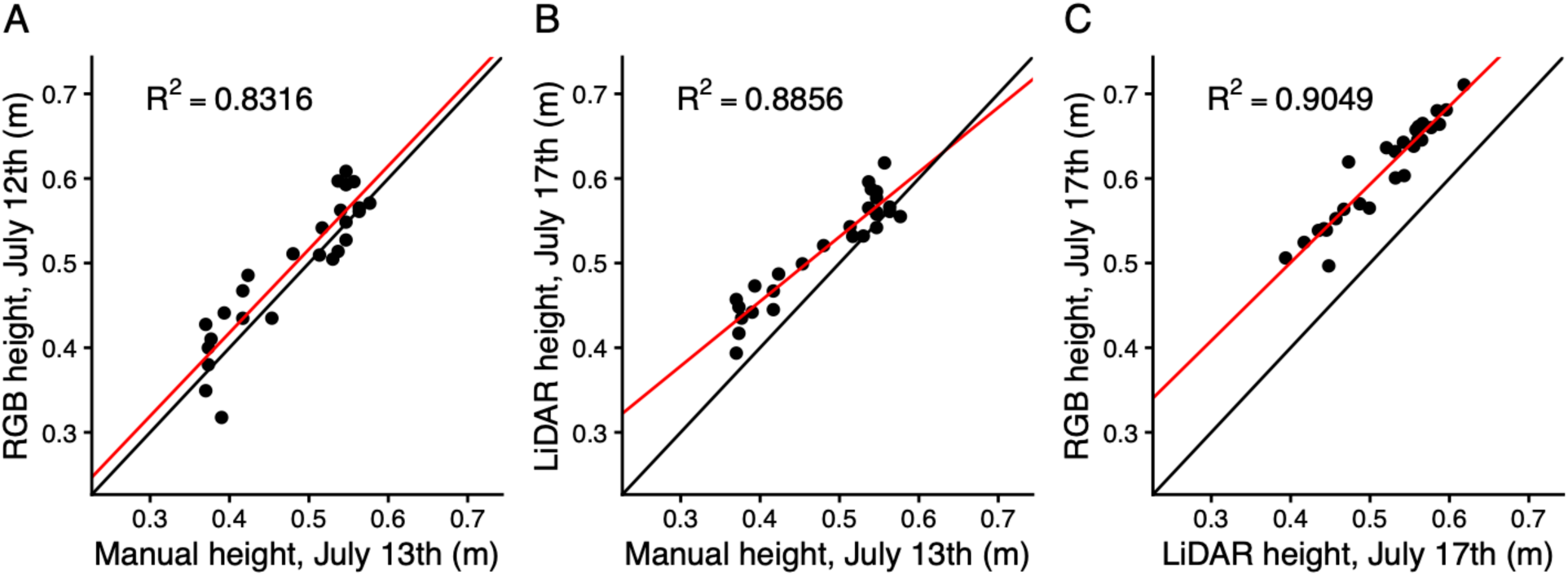
Measurements of height estimated from either RGB images (A) or LiDAR data (B) correlated with manual measurements. RGB-estimated height was also correlated with LiDAR-estimated height (C).

### Height estimated from LiDAR more closely approximates manual measurements

Next, we hypothesized that LiDAR would also correlate to manual stand height measurements. Unlike for RGB images, we did not collect LiDAR within a day of the manual measurements (July 13), so we compared the closest timepoint from a flight four days later (July 17). Despite the intervening days, LiDAR-estimated height predicted manual height measurements accurately (R^2^=0.89, Fig. 3B). RGB-estimated height from images collected on July 12th had a lower correlation to manual measurements compared to LiDAR, despite being closer in time (Fig. 3A).

Importantly, while RGB compared to manual measurements had a slope of 0.985 (95% CI 0.803-1.168), LiDAR slope was significantly different from 1 at 0.764 (95% CI 0.651-0.877) (see intersection of red and black lines in Fig. 3B). The reduced slope of the correlation between LiDAR and manual measurements suggests that either taller plots were underestimated by LiDAR, or that shorter plants grew faster than taller plants in the 4-day span between the manual measurements and LiDAR flight. Which of these hypotheses explains the relationship cannot be determined from this dataset. When comparing RGB and LiDAR measurements captured on the same date (July 17), we found a strong correlation between measurements from both sensor types (R^2^=0.90, Fig. 3C), suggesting they provide similar results despite different methods. The intercept of the correlation between RGB and LiDAR does differ from 0, however (t=4.202, p=0.00295) due to RGB analysis producing larger height estimates. Correlation between rank order is significant (Spearman’s rho = 0.963, p = 2.41e-07) indicating that either method produces measurements with utility for assessing relative height differences between plots.

Because the method for estimating height manually involved averaging the height of a paper laid on top of several places in a plot, we also analyzed the LiDAR data using a method based on this idea (see Methods for more detail). The plot heights as estimated by the LiDAR “paper” method were well correlated with the plot heights estimated by the LiDAR plant and soil threshold subtraction method (R^2^ = 0.986, t = 85.50, p < 2e-16). The slope of the correlation is slightly, but significantly, larger than 1 (95% CI 1.024 - 1.072), indicating that the soil subtraction method produces larger height estimates at larger height values than the “paper” method.

### UAS confirms semidwarf tef lines are shorter than wild-type and more resistant to lodging

Manually collected data confirmed that semidwarf tef lines were more resistant to lodging. We next tested if UAS methods could provide efficient metrics for lodging, rather than the labor-intensive and subjective manual lodging scores. First, we hypothesized that height at a single timepoint after a weather event may be a suitable metric for lodging, and that plots with greater lodging scores would have reduced height. After the storm, wild-type plots had lower average heights compared to semidwarf plots, and remained lower for the remainder of the season (Fig. 4). There was a significant linear correlation (p-value = 6.627e-07) between the manual lodging scores and the UAS measurements of plot height on August 31, which was the last RGB time point and the closest measurement temporally to when lodging was scored.

**Figure 4.**
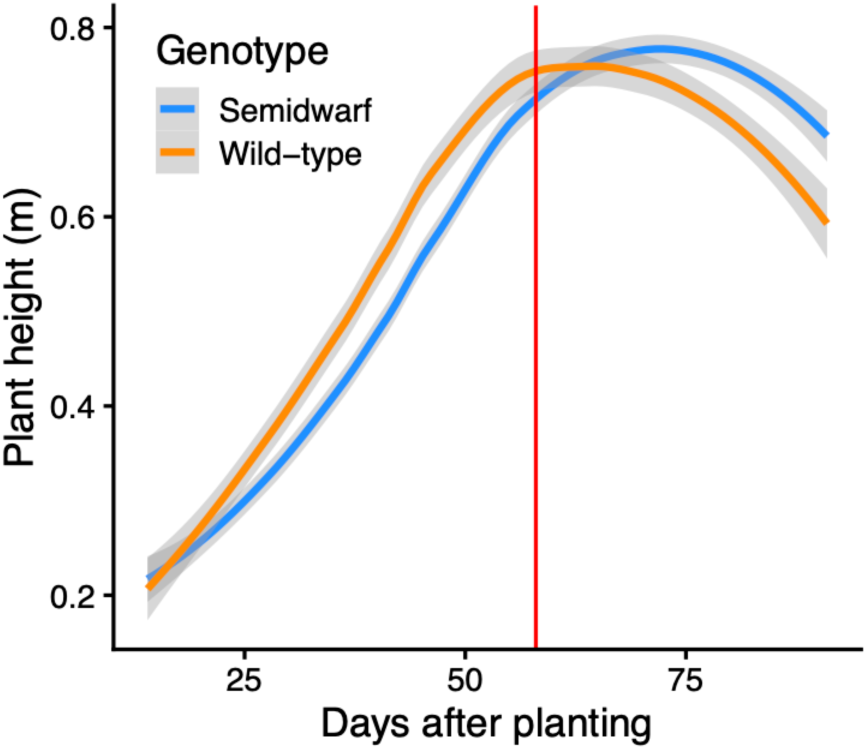
Wild-type plants (orange) were taller than semidwarf lines (blue) early in the season, but lodged more both after a storm event (red vertical line) and progressively towards the end of the season. Colored line represents average of all wild-type or semidwarf lines, respectively.

However, even pre-storm height predicted final manual lodging scores. The plot heights from July 27, the midpoint of the growing season and last timepoint before the storm, had a significant relationship (t-value = 6.439700, p-value < 0.0001) to end-of-season lodging scores using a linear mixed-effects model (Fig. 5A). This relationship indicates that measurements extracted from UAS images at mid-season time points can predict the best performing lines. This suggests that while height after a weather event may be an effective high-throughput measurement of lodging, without additional context of size before an event, it’s not possible to know if the result is due to starting size or the height change.

**Figure 5.**
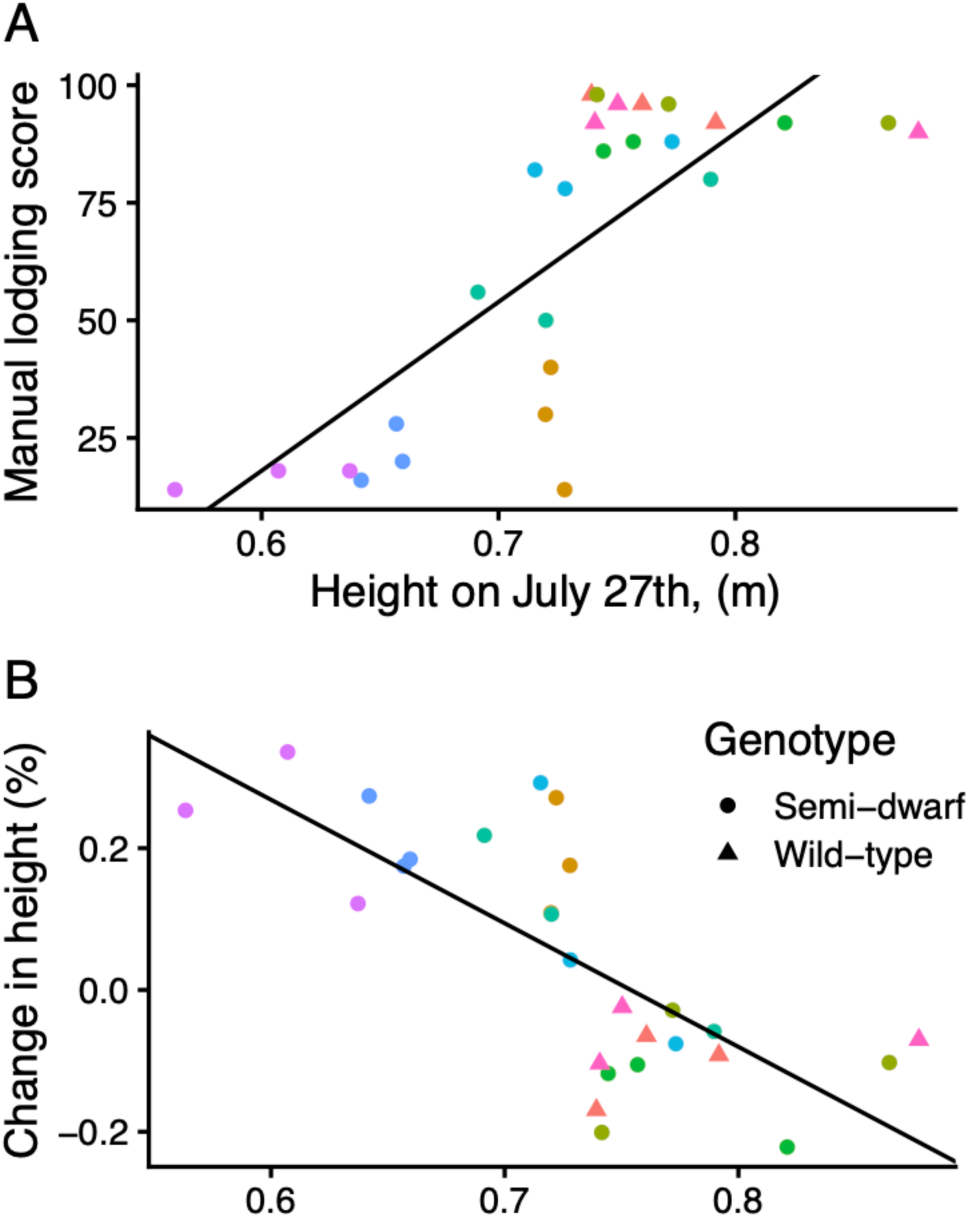
Taller plants experienced a greater decrease in height after a storm (A) and had a higher manual lodging score at the end of the season (B). There was variation in height among semidwarf lines (circles) but the shorter lines were more resistant to lodging. Color represents a replicated semidwarf line (n = 3).

For a more robust UAS measurement of lodging, we measured the change in height using DEMs from RGB images before and after the high-wind event (Figure S1). This change in height was strongly correlated to height before the storm, suggesting taller plants were more likely to lodge (R^2^=0.55, p-value = 1.03e-05; Figure 5B). This measurement of change, rather than absolute height, requires multiple timepoints, but eliminates conflating stand height with lodging, where a short variety may be considered lodged if only compared at one timepoint. We hypothesize that height thresholds might predict lodging (> ∼0.75m) or continued growth (< ∼0.65m), but sample sizes are too small to statistically test for more complex dynamics (Figure 5A). More data could enable explicit fitting of a changepoint model to capture information that could be useful for breeding to a specific height threshold.

## Discussion

Here, we present a method for using UAS sensor data, both RGB and LiDAR, to construct 3D point clouds to estimate height of genetically variable plots of *Eragrostis tef* across a growing season. Both methods for image analysis of point clouds estimate height accurately as compared to manual measurements (Figure 3), but decrease necessary human time, allowing for higher temporal resolution. By imaging at multiple time points through the season, we were able to capture the dynamics of lodging after a high-wind speed storm. Breeding for higher yield in tef requires adopting strategies for reduction in loss from lodging, such as the development of semidwarf varieties. We show that the modified lines used in this study are both shorter and lodge less, and that early season height variance among the lines predicts the severity of lodging.

Our method uses average stand height to compare lines and replicates. While average stand height of a plot is important, the distribution of stand heights within a plot is an important indicator of plant growth and lodging that can be missed when only considering a plot-level average. While a person may be able to make a small number of manual stand height measurements for a plot, UAS imaging measures stand height for every pixel, and thus is often a more accurate representation of the whole plot. In this study, plant height was described as a singular measure, but the level of detail available to extract from this 3D image data will be valuable in understanding growth dynamics, plant health indices, and more. UAS can provide an objective and data-rich approach to describing plot height and offers a higher throughput approach to quantification in lodging studies in the field.

We found that LiDAR and RGB analyses both produced similar results for stand height. While LiDAR provided a higher correlation to manual measurements (Figure 3), the data sizes were larger, analysis was more complex, and the equipment was more costly.

RGB images also provide additional data on plot color that could be analyzed with additional calibration during photogrammetry for other research questions, such as plant health, flowering time, and senescence. We recommend researchers consider the objectives of the experiment and the expected height differences of their control and test lines when determining which sensor is appropriate to achieve the experimental goals while keeping costs and analysis time low. For example, more subtle differences between control and test varieties may require the higher accuracy afforded by LiDAR to detect differences.

The height analysis from DEMs described above does not require elevation values to be georectified to an absolute coordinate system, since height is determined by the difference in elevation between plants and soil within a plot. This approach may be less accurate when bare soil is not visible, such as when there is total canopy coverage, but depending on the shape of plot height distributions, optimizing for a threshold may still work. Residue (such as from no-till management), cover crops in biculture, and weeds would also impact the appropriate approach to using UAS for plot heights. We also found that the specific plot boundaries (i.e. how much soil area is included) can affect the appropriate choice for soil and plant thresholds for the pixel distributions, which are used to calculate average plot height. As such, we recommend visually inspecting the shape of pixel height distributions (such as in Figure 2A) prior to running this analysis to help choose appropriate threshold values, as was performed here (see Methods). Even so, this method can produce variability due to stochasticity in plot boundaries, which should be considered when interpreting results.

Importantly, the occurrence of the storm during this experiment was unintentional, and obscured possible differences in lodging later in the season that can occur during grain filling. It is possible that the effect of height is consistent across both lodging from wind or rain and lodging from grain weight, but that cannot be determined in this study. Instead, we focus on the ability of 3D point clouds to aid in both the tracking of height changes through a season and on the prediction of possible yield loss from lodging. Because of the correlation with both manual height measurements at a single time point and the increased time resolution afforded, we conclude that UAS image analysis could provide a benefit to tef breeding strategies that target resistance to lodging.

## Acknowledgements

UAV data were collected and provided by Remote Sensing Lab at Saint Louis University as part of a Taylor Geospatial Institute Block Grant to the Donald Danforth Plant Science Center. We acknowledge the use of Data to Science (D2S, https://d2s.org) platform, an open-source project developed by Geospatial Data Science Lab (https://gdsl.org) at Purdue University. We thank the Phenotyping Core Facility (RRID:SCR_019049), particularly Joseph Duenwald for ground control point maintenance, and the Field Research Site at the Donald Danforth Plant Science Center for plant care. The developers of PlantCV-Geospatial thank Sam Taylor and Jalissa Pirro for help and guidance.

## Author Contributions

KMM, GB, and NF designed experiments. GB developed tef lines, planted, and performed manual measurements of height and lodging score. KEB, HS, ML, and DS performed data analysis. KEB, HS, and KMM wrote the article with contributions from all authors.

## Data Availability

Code and data associated with this manuscript are available on GitHub (https://github.com/danforthcenter/teff-manuscript).

---

**Supplemental Figure S1:**
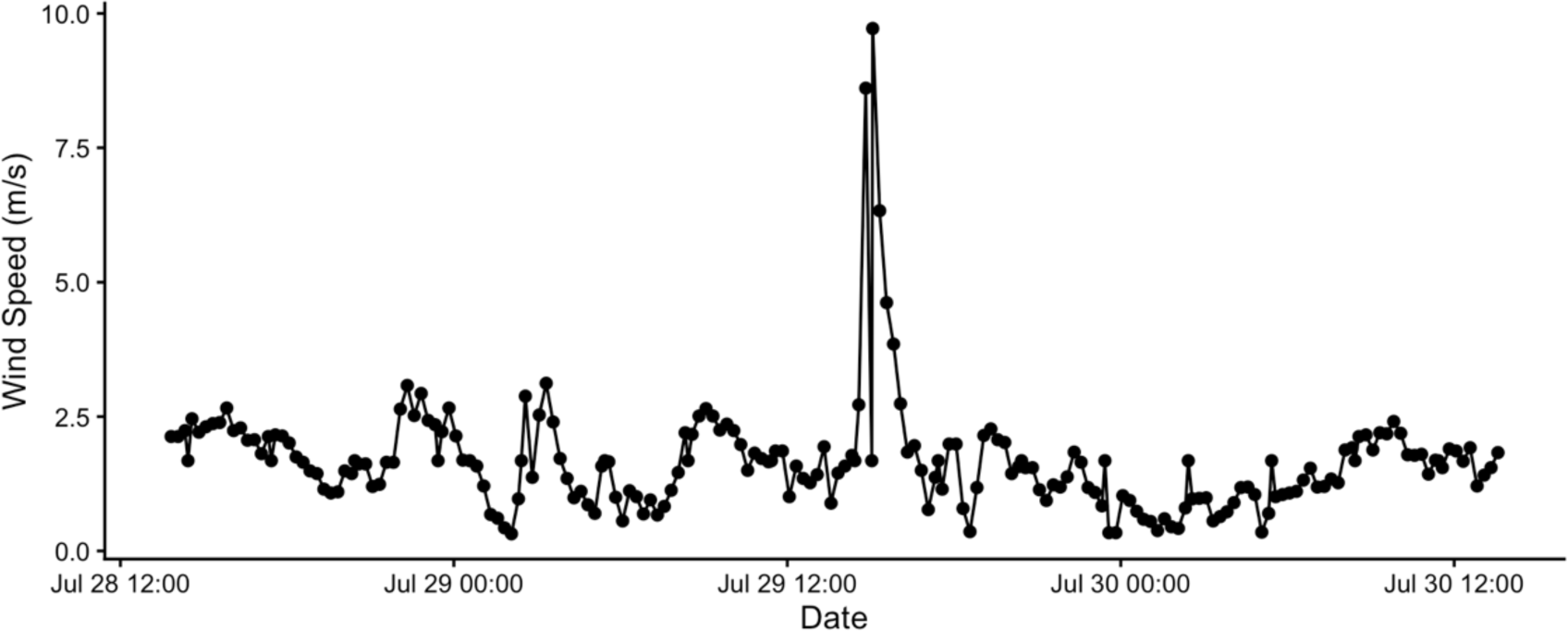
Wind speed measured over a 48h period during which a storm occurred in the vicinity of the field plots.

